# Preferential Pharmacological Inhibition of Nav1.6, but not Nav1.1, Abolishes Epileptiform Activity Induced by 4-AP in Mouse Cortical Slices

**DOI:** 10.1101/2020.05.29.124693

**Authors:** Reesha R. Patel, Xingjie Ping, Shaun R. Patel, Jeff S. McDermott, Jeffrey L. Krajewski, Xian Xuan Chi, Eric S. Nisenbaum, Xiaoming Jin, Theodore R. Cummins

**Author notes:** CORRESPONDING AUTHOR: Theodore R. Cummins, 723 W. Michigan St, Indianapolis, IN 46202.

## Abstract

Brain isoforms of voltage-gated sodium channels (VGSCs) have distinct cellular and subcellular expression patterns as well as functional roles that are critical for normal physiology as aberrations in their expression or activity lead to pathophysiological conditions. In this study, we asked how inhibition of select isoforms of VGSCs alters epileptiform activity to further parse out the roles of VGSCs in the brain. We first determined the relative selectivity of recently discovered small molecule, aryl sulfonamide, inhibitors (ICA-121431 and Compound 801) against Nav1.1, Nav1.2, and Nav1.6 activity using whole-cell patch clamp recordings obtained from HEK293 cells. To test the effects of these inhibitors on epileptiform activity, we obtained multielectrode array (MEA) recordings from mouse cortical slices in the presence of 4-aminopyridine (4-AP) to induce epileptiform activity. We found that the ICA-121431 and Compound 801 compounds are relatively selective for Nav1.1 and Nav1.6, respectively. From the MEA recordings, we found that inhibition of Nav1.6 and Nav1.2 with 500nM of the Compound 801 compound completely abolishes ictal local field potentials induced by 4-AP, whereas inhibition of Nav1.1 with 500nM of the ICA-121431 compound had minimal effect on epileptiform activity induced by 4-AP. Due to the prominent expression of Nav1.1 in inhibitory neurons, we asked whether inhibition of Nav1.1 alone alters activity. We found that, indeed, inhibition of Nav1.1 with the ICA-121431 compound increased basal activity in the absence of 4-AP. These findings expand our current understanding of the roles of VGSC isoforms in the brain and suggest that selective targeting of Nav1.6 may be a more efficacious treatment strategy for epileptic syndromes.

## INTRODUCTION

Voltage-gated sodium channels (VGSCs) are critical for neuronal excitability as they are responsible for the initiation and propagation of action potentials. They comprise of a large α subunit (Nav1.1 – Nav1.9) that can associate covalently or non-covalently with several auxiliary proteins including one or more β subunits (Navβ1 – Navβ4) [1]. Therefore, these channels function as an intricate channel complex that is highly regulated. Structurally, these channels consist of four homologous domains each with six transmembrane segments (S1-S6) wherein the S1-S4 segments make up the voltage-sensing modules while the S5-S6 segments make up the central pore of the channel. Conformational changes due to movement of the voltage-sensing modules within the protein lead to state transitions of the channel. At hyperpolarized, resting membrane potentials, channels are primarily in a closed (non-conducting) state and upon membrane depolarization they transition to an open state, conducting an inward sodium current, and within milliseconds after opening they typically enter an inactivated state (non-conducting). The inward, depolarizing current generated by these channels underlies the rising phase of the action potential, making VGSCs essential for the electrical signals by which neurons communicate.

There are four major VGSCs isoforms expressed in the central nervous system: Nav1.1, Nav1.2, Nav1.3, and Nav1.6 [1]. These brain isoforms of VGSCs have distinct cellular (i.e. neuronal populations) as well as subcellular (i.e. neuronal compartments) expression patterns. Nav1.1 is expressed in different neuronal compartments including: soma, dendrites, nodes of Ranvier as well as the axon initial segment [2-5]. Although much of our understanding comes from rodents, it is believed that distributions are generally similar in humans. Nav1.1 is predominantly expressed in parvalbumin and somatostatin positive GABAergic neurons [6-12], although it has been observed in some excitatory neuronal populations [13]. Nav1.2 is highly expressed in the proximal axon initial segment, unmyelinated axons and nerve terminals [5, 14-18]. Specifically, Nav1.2 expression has been observed in the proximal axon initial segment of excitatory, pyramidal neurons [19, 20]. Some reports indicate Nav1.2 is also expressed in somatostatin positive GABAergic neurons [21], but this has been questioned [12]. While Nav1.3 mRNA has been detected during prenatal development, its expression diminishes during early postnatal days and is lost in the adult rodent brain [5, 22]. Interestingly, Nav1.3 is widely detected in adult human brains [4]. Nav1.6 is ubiquitously expressed in the brain. In particular, Nav1.6 expression is dense in the axon initial segment and nodes of Ranvier of many neuronal populations [14, 15, 23-25]. Each isoforms’ function within these different regions is highly specialized as evidenced by disruption of their activity or expression leading to pathological conditions and/or premature lethality [26-28]

It is not surprising from their critical role in excitability that VGSCs have been implicated in epilepsy. The most convincing evidence for this are genetic epilepsies due to mutations in VGSCs. Indeed, mutations have been identified in all brain isoforms of VGSCs and result in distinct epileptic syndromes [29-32]. Much insight into the role of VGSCs in epilepsy has been gleaned from studying animal models of these genetic epilepsies. To date, the leading hypothesis for the mechanism underlying Nav1.1 mutations is the decreased activity of inhibitory interneurons while little to no effect on excitatory neurons, due to the prominent expression of Nav1.1 in inhibitory neurons, leading to an overall decrease in inhibitory tone [7, 11, 27, 33, 34]. Recently, the first epilepsy-associated mutation in Nav1.6 (N1768D) was identified and found to cause gain-of-function in heterologous expression systems – increasing both Nav1.6 activity and excitability of hippocampal neurons [35]. Heterozygous *Scn8a*^N1768D/+^ knock-in mice exhibit seizures and sudden unexpected death in epilepsy demonstrating the causal role of this mutation [36]. These findings support the primary hypothesized mechanism by which epilepsy-associated mutations in Nav1.6 lead to epilepsy is due to increased activity of Nav1.6 and a consequent increase in overall neuronal excitability due to the ubiquitous expression of Nav1.6 in the brain. Interestingly, seizure thresholds and premature lethality of Nav1.1 heterozygous mice, a model for severe myoclonic epilepsy in infancy (SMEI), can be restored and rescued to wildtype levels by essentially eliminating a single Nav1.6 allele, suggesting that a reduction in Nav1.6 activity may decrease seizure generation [37]. Indeed, heterozygous Nav1.6 mice have increased thresholds to chemically and electrically induced seizures [37, 38]. Moreover, Nav1.6 channel expression and activity is increased in kindling models of epilepsy as well as genetic epilepsies such as those due to Celf4 deficiency [39-42]. Overall, these findings suggest that broadly inhibiting Nav1.1 activity could have adverse effects, while selectively inhibiting Nav1.6 may be more beneficial for the treatment of epilepsy syndromes.

VGSCs are the target for many drugs used clinically including: local anesthetics, anti-arrhythmias and anti-epileptics (AEDs). The primary mechanism of action for commonly used anti-epileptics such as phenytoin, carbamazepine, lamotrigine and others is to inhibit VGSC activity. These drugs bind to the pore of the channel with higher affinity for channels in an open or inactivated state, thus endowing them with a use-dependence property [43]. This region of the channel is highly conserved among VGSC isoforms, therefore making classic anti-epileptics relatively nonselective. While classic AEDs can help control seizures in some patients, approximately 20 to 40% of epilepsy patients are refractory to treatment. Therefore, there is a need to develop novel AEDs. From animal model studies, it seems plausible that selectively targeting specific isoforms of VGSCs may be a more efficacious strategy for the treatment of epilepsy syndromes.

Here we focused on the contribution of Nav1.1, Nav1.2 and Nav1.6 to epileptic activity due to their distinct expression pattern and availability of selective small molecule inhibitors of these channel isoforms. We first determined the selectivity of the Compound 801 (801 cmpd) and ICA-121431 (ICA cmpd) for Nav1.1, Nav1.2 and Nav1.6 using whole-cell patch clamp recordings in HEK293T cells. We then asked whether preferential inhibition of Nav1.1 or Nav1.6 by ICA and 801 cmpds, respectively, would have differential effects on epileptic activity. To address this question we used a 4-aminopyradine (4-AP) model of epileptiform activity, previously characterized by others [44, 45], and multielectrode array (MEA) recordings from mouse cortical slices. We found that preferential inhibition of Nav1.6 abolishes epileptiform activity, while inhibition of Nav1.1 did not. This led us to ask if inhibition of Nav1.1 alone could induce synchronized activity. We found that partial inhibition of Nav1.1 activity in mouse cortex can increase basal activity. Our findings suggest that selective targeting of VGSC isoforms, in contrast to current AEDs that broadly inhibit VGSCs, may be a more efficacious treatment strategy for epileptic syndromes.

## MATERIALS and METHODS

### cDNA Constructs

Optimized human constructs for Nav1.1, Nav1.2 and Nav1.6 were designed in-house and purchased from Genscript (Piscataway, NJ). cDNA constructs for wildtype Nav1.1, Nav1.2 and Nav1.6 channels encode for amino acid sequences corresponding to the accession numbers BAC21102.1, NP_001035232.1 and NP_055006.1 in the NCBI database, respectively.

### Cell Cultures and Transfections

The use of HEK293T cells [46] was approved by the Institutional Biosafety Committee and followed the ethical guidelines for the National Institutes of Health for the use of human-derived cell lines. HEK293T cells were grown under standard tissue culture conditions (5% CO2; 37°C) with DMEM supplemented with 10% fetal bovine serum. HEK293T cells were transiently transfected using the calcium phosphate precipitation method. Briefly, calcium phosphate-DNA mixture (4.5 μg channel construct and 0.5 μg EGFP) was added to cells in serum-free media for 4-5 hours after which it was replaced with normal media. 12-24 hours post-transfection, cells were split onto laminin-coated glass coverslips. Cells were identified by expression of EGFP using a fluorescent microscope and whole-cell patch clamp recordings were obtained 36-72 hours post-transfection. Stable HEK cell lines expressing mNav1.1, mNav1.2 and mNav1.6 were used to determine the pharmacological sensitivity of mouse sodium channels.

### Chemicals and Solutions

ICA-121431 (ICA cmpd) and 4-aminopyridine (4-AP) was obtained from Sigma Aldrich Co. (St. Lousi, MO). Compound 801 (801 cmpd) was a generous gift from Eli Lilly and Co. (Indianapolis, IN) and was identified in Icagen/Pfizer patent WO2010079443 and synthesized at Lilly Laboratories (Indianapolis, IN). ICA and 801 cmpds were dissolved in 1-methyl-2-pyrrolidinone (MPL) to a stock concentration of 10mM and stored at 4°C. 4-AP was also dissolved in MPL to a stock concentration of 100mM and stored at 4°C. All drugs were diluted to desired concentration in artificial cerebrospinal fluid (ACSF) or extracellular patch-clamp solution just prior to use.

### Whole-Cell Patch Clamp Recordings

Whole-cell patch clamp recordings were obtained at room temperature (∼23°C) using a HEKA EPC-10 amplifier and the Pulse program (v 8.80, HEKA Electronic, Germany) was used for data acquisition. Electrodes were fabricated from 1.7 mm capillary glass and fire-polished to a resistance of 0.9-1.3 MΩ using a Sutter P-97 puller (Sutter Instrument Company, Novato, CA). All voltage protocols were started 5 minutes after obtaining a gigaΩ seal and entering the whole-cell configuration, which controlled for time-dependent shifts in channel properties. Voltage errors were minimized to less than 5 mV using series resistance compensation and passive leak currents were cancelled by P/-5 subtraction. The bath solution contained in (mM): 140 NaCl, 1 MgCl2, 3 KCl, 1 CaCl2, and 10 Hepes, adjusted to a pH of 7.30 with NaOH. The pipette solution contained in (mM): 140 CsF, 10 NaCl, 1.1 EGTA, and 10 Hepes, adjusted to a pH of 7.30 with CsOH. Recordings were made in the presence of extracellular solution containing the drug. Each coverslip was recorded from for up to one and half hours before discarding.

### Cortical Slice Preparation

Male and female C57BL/6 mice from postnatal day 11 to 16 were anesthetized with ketamine and decapitated. Brains were quickly removed and placed in chilled, oxygenated dissecting solution containing (in mM): 111 choline-Cl, 2.5 KCl, 1.25 NaH2PO4, 10 MgSO4, 0.5 CaCl2, 26 NaHCO3, and 10 D-glucose (pH 7.4 and ∼305mOsm). Brains were bisected along the sagittal plane, leaving the left and right hemispheres intact and coronal cortical slices were cut to a thickness of 300μm using a vibratome (Leica VT1200, Buffalo Grove, IL). After cutting, slices were incubated in oxygenated artificial cerebrospinal fluid (ACSF) for at least 1 hour at room temperature before being used for recording. ACSF solution contained (in mM): 126 NaCl, 2.5 KCl, 1.25 NaH2PO4, 2 MgSO4, 2 CaCl2, 26 NaHCO3 and 10 D-Glucose (pH 7.4 and ∼315mOsms).

### Multielectrode Array Recordings

MEA recordings were performed similarly to methods previously described [47]. Briefly, multielectrode array probes were coated with 0.1% polyethylenimine in boric solution for approximately three hours and washed with deionized water. Slices were placed by eye in a manner that neocortical layers I-V covered the array (Figure 2). Slices were maintained at 37°C and continuously perfused with oxygenated ACSF containing drugs. Experiments testing drug effects on 100μM 4-AP induced hyper-excitability were conducted as follows: one hour of baseline (100μM 4-AP only) recording followed by thirty minutes of treatment (100μM 4-AP and drug of interest) recording and lastly washout (100μM 4-AP only) to confirm slice viability. For experiments testing the effects of drugs on basal activity, a forty minute recording of activity in the presence of normal ACSF or drug alone was followed by a 20 minute recording in the presence of 100μM 4-AP, to assure slice viability, was obtained. Activity was sampled from all sixty electrodes at a sampling rate of 1kHz and amplified using MC_Rack software from Multichannel Systems (Reutlingen, Germany). Data was stored for further offline analysis.

**Figure 1.**
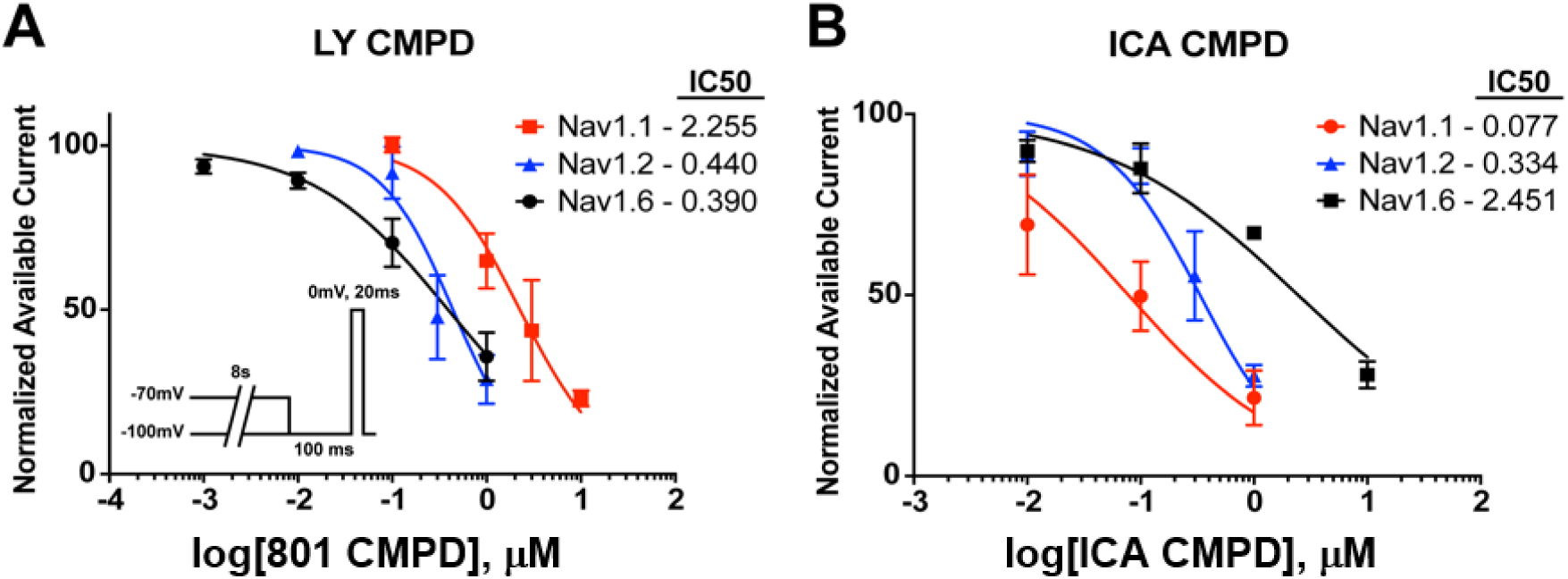
Concentration response curves for 801 and ICA compounds on human voltage-gated sodium channels. A, Concentration response curves for human Nav1.1 (red squares), Nav1.2 (blue triangles) and Nav1.6 (black circles) with the 801 compound. B, Concentration response curves for hNav1.1 (red circles), hNav1.2 (blue triangles) and hNav1.6 (black squares) with the ICA compound. To assess current inhibition, a conditioning step to -70mV for 8s, allowing drug to bind the channel, followed by a recovery step to -100mV for 100ms and subsequently a final step to 0mV for 20ms was applied to measure available current (*inset*). The available current was normalized to the maximum current measured with a similar voltage protocol with a conditioning step to -100mV. Each data point represents an *n* = 2-8 of separate experimental cells.

**Figure 2.**
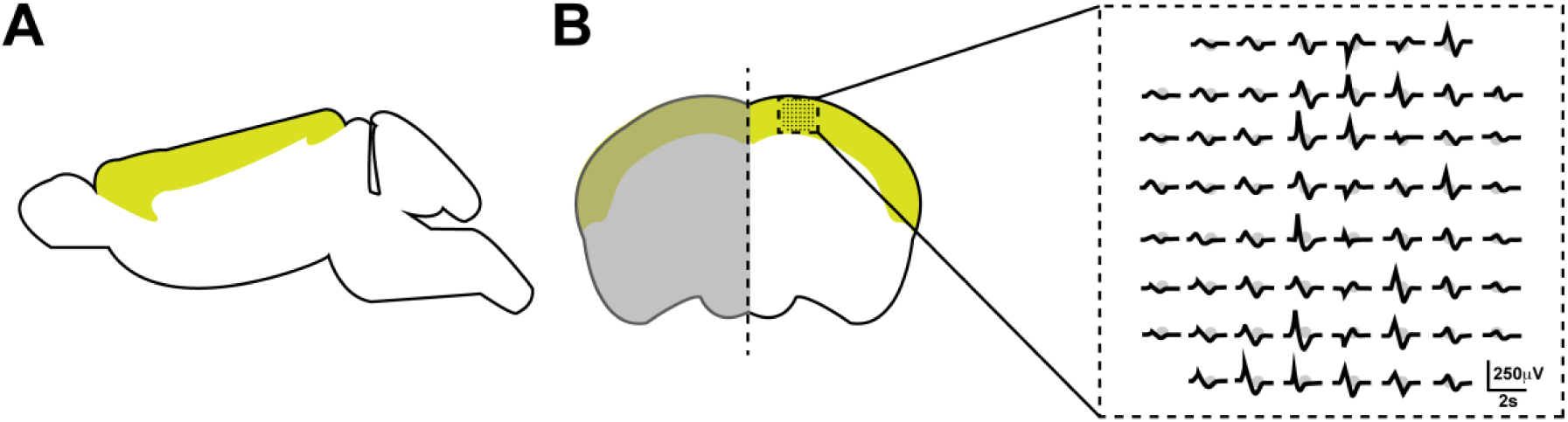
Multielectrode array recordings. A, Sagittal view of mouse brain highlighting cortex region in green. B, Coronal brain slice depicting the approximate placement of the multielectrode array in the neocortical layers I-V. An illustration of the arrangement of the sixty electrodes, which were 30μm in diameter and height and spaced 200μm apart, overlaid with typical recorded activity, can be seen in the dotted box.

### Data Analysis

Whole-cell voltage clamp data are represented as the mean ± the standard error and *n* reflects the number of separate experimental cells. Concentration response curves were fit with the Hill logistic equation: y = 100/(1+10^((LogIC_50_-x) * HillSlope))) to obtain IC50 values using Graphpad Prism (La Jolla, CA).

Analysis of multielectrode array data was done using an in-house generated MATLAB script. A twenty-minute window in the middle of each recording was used for analysis of activity. Data was filtered using a second order butterworth filter with a cutoff frequency of 15Hz to get rid of high frequency noise and isolate slow local field potentials (LFPs). The script returns peak values above or below a threshold set to at least six times the standard deviation of the noise (after filtering) measured for 20 seconds during which no LFPs occur for each recording. The top ten most active electrodes, as determined by the number of events, during the baseline recording from each slice were used for further analysis. Peak amplitude was calculated as the average of all peak values from a single electrode and subsequently averaged across the top ten electrodes. Valley amplitudes were calculated in a similar manner. The duration of the LFPs was measured from the mean waveform from each electrode. Duration values were then averaged across the top ten electrodes. The mean waveform was obtained by plotting LFPs aligned by their peak values. Only LFPs of the most common type, which always included ictal events, were used for measurement of duration. MATLAB and Graphpad Prism (La Jolla, CA) were used to make graphs. Black data points represent the mean value from each slice, while the mean of all slices in the group are represented in red data points. The sample number, *n*, signifies the number of separate experimental slices. Statistical significance was determined using either a paired, when appropriate, or unpaired t-test.

## RESULTS

### Preferential inhibition of Nav1.1 and Nav1.6 by ICA and LY compounds, respectively

Recently, it was discovered that small molecule, aryl sulfonamide, inhibitors can selectively target VGSC isoforms [48, 49]. In this study, we examined the effects of the ICA-121431 compound (ICA cmpd) and a derivative of the previously described PF-04856262, the Compound 801 (801 cmpd) [49]. The selectivity of these molecules has been attributed to specific extracellularly accessible amino acid residues of the voltage-sensing module of domain IV. Based on the conservation of these residues in the different isoforms, we hypothesized that these compounds would differentially inhibit Nav1.1 and Nav1.6. Indeed, the ICA cmpd has previously been found to inhibit Nav1.3 and Nav1.1 (IC50 = ∼10-20nM) to a greater extent than Nav1.2 (IC50 = ∼250nM) and much more potently than Nav1.6 (IC50 > 10μM) [49]. To test the selectivity of these compounds on Nav1.1, Nav1.2, and Nav1.6 activity, we obtained whole-cell patch clamp recordings from HEK293 cells stably expressing mouse voltage-gated sodium channels (Table 1). Relative sensitivities of mouse Nav1.1, Nav1.2 and Nav1.6 channels to the compounds was determined with test pulses to -10 mV. Mouse Nav1.1 was significantly less sensitive to the 801 cmpd than the ICA cmpd. By contrast, mouse Nav1.2 was significantly more sensitive to the 801 cmpd than the ICA cmpd. Mouse Nav1.6 was also significantly more sensitive to the 801 cmpd than the ICA cmpd.

**Table 1.** IC50s for compounds on mouse voltage-gated sodium channels.

|  | mNav1.1 | mNav1.2 | mNav1.6 |
| --- | --- | --- | --- |
| 801 cmpd | $261.1 \pm 13.3\text{ nM}$, n=9* | $4.2 \pm 2.0\text{ nM}$, n=12 | $1.6 \pm 0.9\text{ nM}$, n=16 |
| ICA cmpd | $10.8 \pm 1.0\text{ nM}$, n=13* | $553.4 \pm 60.6\text{ nM}$, n=19# | $3.7 \pm 1.1\text{ }\mu\text{M}$, n=13 |

\* Significantly different from mNav1.2 and mNav1.6. #Significantly different from mNav1.6. ANOVA with Fisher's LSD post-hoc test, $p < 0.05$ .

We also examined the selectivity of these compounds on human sodium channel activity with whole-cell patch clamp recordings from HEK293T cells transiently transfected with human Nav1.1, Nav1.2, and Nav1.6 channels. We used a voltage command protocol in which we applied an initial conditioning pulse to -100mV or - 70mV, to determine the maximum and inhibited current, respectively, for 8 seconds followed by a recovery pulse to -100mV for 100ms and a final test pulse to 0mV for 20ms (Figure 1A *inset)*.We generated concentration response curves for each compound and human isoform to determine the concentration at which fifty percent of activity was inhibited (IC50) (Figure 1). We found that the 801 cmpd more potently inhibits hNav1.6 (IC50 = 0.39μM) and hNav1.2 (IC50 = 0.44μM) compared to hNav1.1 (IC50 = 2.26μM) (Figure 1A). In contrast, the ICA cmpd is selective for hNav1.1 (IC50 = 0.08μM) over hNav1.2 (IC50 = 0.33μM) and hNav1.6 (IC50 = 2.25μM) (Figure 1B). Under these conditions, the compounds showed less potency compared to the potency observed with the mouse isoforms. This can be explained in part by the use of a different voltage command protocol to measure sensitivity. With the human isoforms we used an 8 second conditioning pulse to -70mV to mimic a standard physiological resting membrane potentials and thus obtain a physiologically relevant measure of inhibition and relative selectivity. Importantly, the relative order of sensitivity is the same for the mouse and human isoforms. Thus the differential selectivity of these compounds, especially with respect to the mouse channels, gave us the opportunity to ask how different VGSC isoforms contribute to epileptic activity.

### Preferential inhibition of Nav1.2 and Nav1.6, but not Nav1.1, abolishes epileptiform activity

We used multielectrode array recordings from mouse cortex to study the effects of these selective, small molecule inhibitors on 4-AP induced epileptiform activity (Figure 2). We first examined the effects of continuous application of 100μM 4-AP for 1.5 hours on cortical activity to determine if activity changes over time. Representative traces of activity recorded during the first hour which is referred to as the baseline recording (100μM 4-AP) and the next thirty minutes which is referred to as the treatment recording (100μM 4-AP - Control) can be seen in Figure 3A. We found that there was no significant change in the number of ictal LFP events between baseline and treatment recordings. Baseline activity induced by 4-AP was variable between slices; therefore we performed paired experiments and show the mean value from each slice (black data points) as well as the overall mean for each group (red data points) (Figure 3). As seen in Figure 3B, the ictal like LFP events were biphasic with an upstroke to a peak followed by downstroke to a valley. The peak and valley amplitudes as well as the duration of the LFPs remained the same over time (Figure 3C-F). This allowed us to directly measure drug effects on 4-AP induced activity without having to compensate for run –up or –down effects of the model.

**Figure 3.**
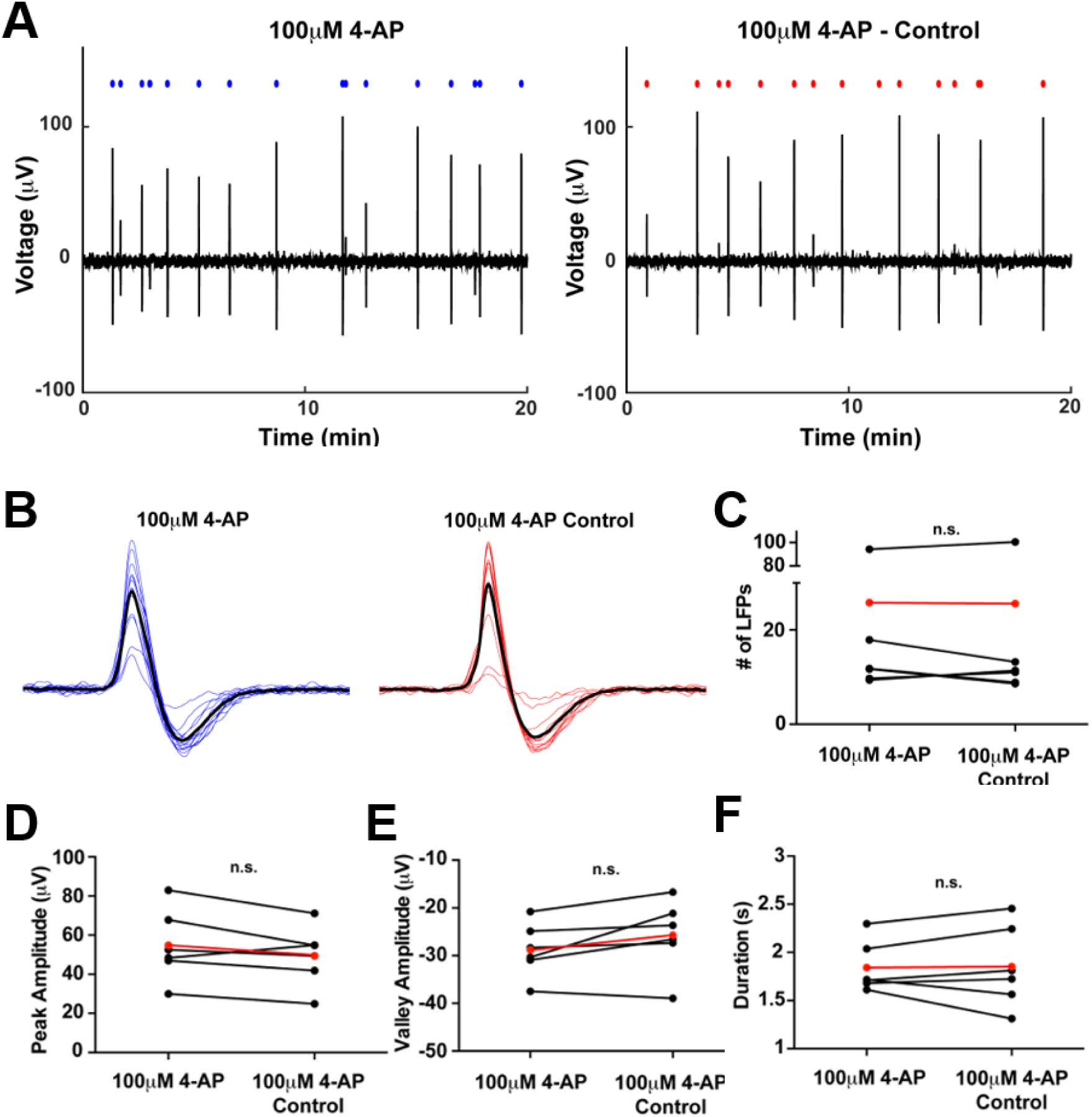
100μM 4-AP induced hyper-excitability in mouse cortical brain slices. A, Representative twenty minute traces of LFP activity from baseline recording (left) and corresponding treatment recording (right). The blue and red circles above traces depict the location of each LFP detected. B, Alignment of all LFPs from a single electrode during baseline recording (blue) and treatment recording (red). The bolded black traces represent the mean LFP waveform from which the duration was measured. C, Average number of LPFs detected from baseline and corresponding treatment recordings (*n* = 6). Data in red represent the mean across all slices. D and E, Average peak and valley amplitude calculated from baseline and treatment recordings. F, Average duration of LFPs recorded during baseline and corresponding treatment groups. n.s., not significant; paired t-test.

We next tested the effects of the 500nM 801 cmpd on 4-AP induced epileptiform activity, which preferentially inhibits Nav1.6 and Nav1.2, by approximately fifty percent according to the IC50 values we observed, while having little to no effect on Nav1.1 or Nav1.3 channel activity at this concentration. We recorded baseline activity for one hour with 100μM 4-AP to induce epileptiform activity and subsequently recorded the treatment condition consisting of 100μM 4-AP with 500nM 801 cmpd for thirty minute. We found that 500nM 801 cmpd completely abolishes ictal LFPs (Figure 4A). We did not observe ictal LFPs in any recordings with the 801 cmpd. The number of LFP events detected was significantly (p<0.05; paired t-test; *n* = 5) reduced in the presence of the 801 cmpd (Figure 4C). Some slices still displayed activity, but of much smaller amplitude (Figure 4D, E). This activity did not exhibit the typical ictal LFP waveform. To confirm slice viability and intact attachment of the slice to electrodes after treatment perfusion, we recorded a washout recording with only 100μM 4-AP that demonstrated activity could be recovered to some extent (Figure 4A *inset*). These data show that partial inhibition of Nav1.6 and Nav1.2 by the 801 cmpd selectively abolishes epileptiform activity induced by 4-AP but does not completely inhibit slice activity.

**Figure 4.**
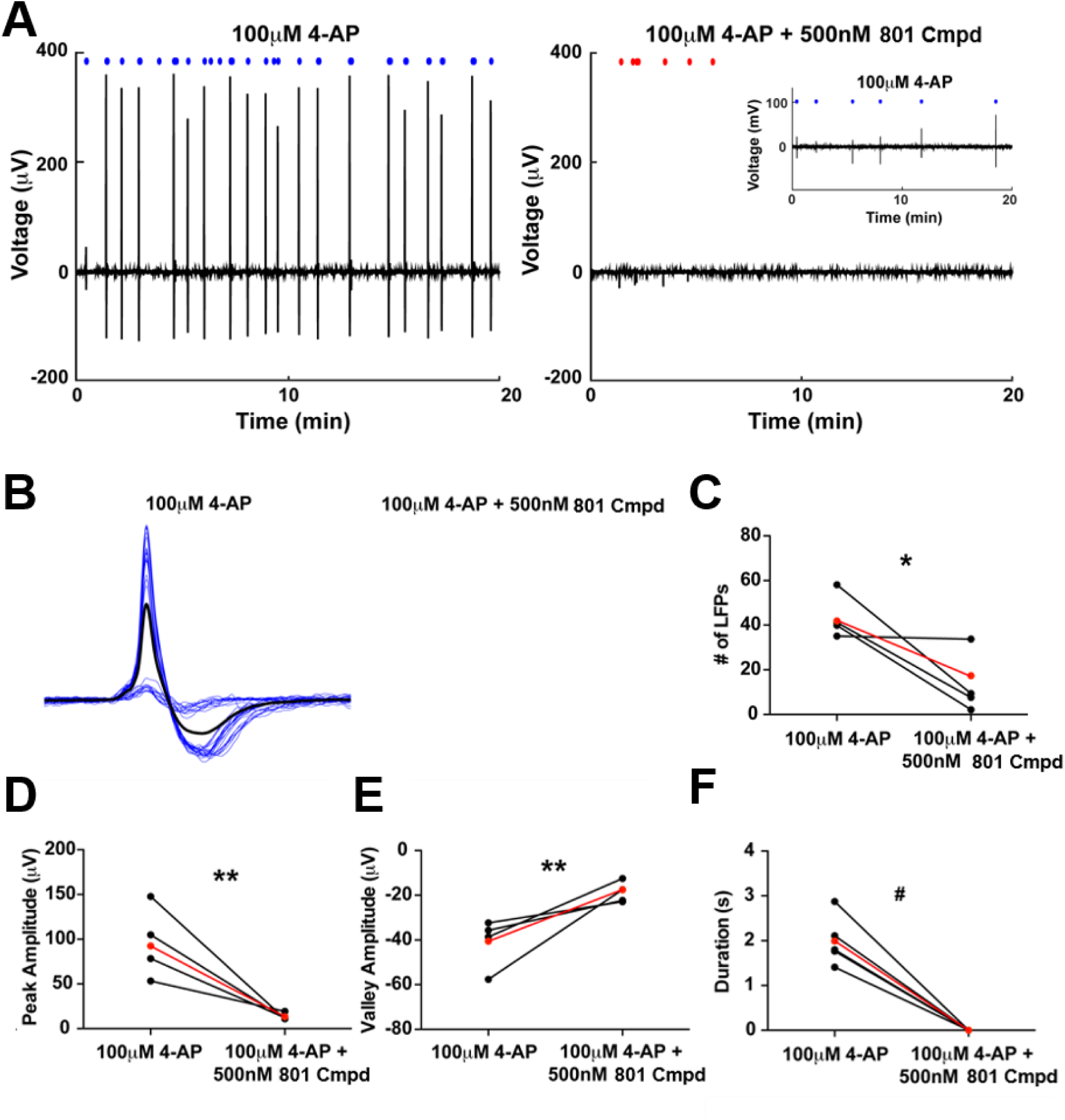
Effects of 500nM 801 cmpd on 4-AP induced hyper-excitability in mouse cortical brain slices. A, Representative twenty minute traces of LFP activity from baseline recording with 100μM 4-AP (left) and corresponding treatment recording with 100μM 4-AP and 500nM 801 cmpd (right). Twenty minute trace of corresponding washout recording with 100μM 4-AP only (*inset*). The blue and red circles above traces depict the location of each LFP detected. B, Alignment of all LFPs from a single electrode during baseline recording (blue). No ictal LFPs were detected during treatment recording. The bolded black trace represents the mean LFP waveform from which the duration was measured. C, Average number of LPFs detected from baseline and corresponding treatment recordings (*n* = 5). Data in red represent the mean across all slices. D and E, Average peak and valley amplitude calculated from baseline and treatment recordings. F, Average duration of LFPs recorded during baseline and corresponding treatment groups. *p<0.05, **p<0.01, ^#^p<0.001; paired t-test.

We further investigated the effects of the 125nM ICA cmpd, which preferentially inhibits Nav1.1 and Nav1.3 while having little to no effect on Nav1.2 and Nav1.6 at this concentration, on 4-AP induced activity. Representative traces of baseline and treatment recordings can be seen in Figure 5A. We found no siginficant effects of 125nM ICA cmpd on the number of ictal LFPs or peak or valley amplitudes of the LFPs (Figure 5C-E). Alinged LFPs from a single representative electorde during baseline and treatment recordings can be seen in Figure 5B. Interestingly, there was a slight but significant (p<0.01; paired t-test; *n* = 5) reduction in LFP duration in the presence of 125nM ICA cmpd. These data suggest that inhibition of Nav1.1 and Nav1.3 has minimal effects on 4-AP induced epileptiform activity.

**Figure 5.**
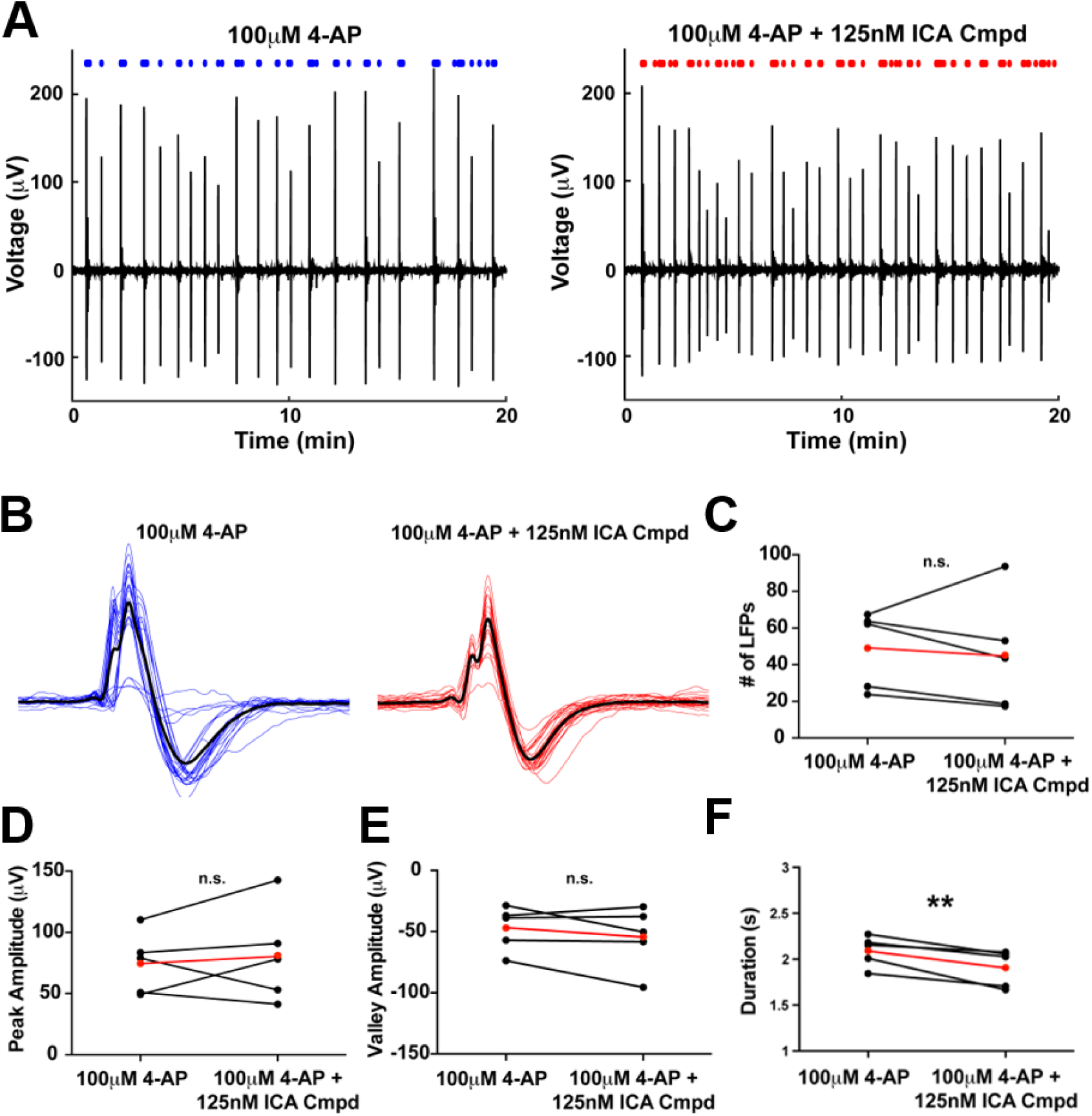
Effects of 125nM ICA compound on 4-AP induced hyperexcitability in mouse cortical brain slices. A, Representative twenty minute traces of LFP activity from baseline recording with 100μM 4-AP (left) and corresponding treatment recording with 100μM 4-AP and 125nM ICA cmpd (right). The blue and red circles above traces depict the location of each LFP detected. B, Alignment of all LFPs from a single electrode during baseline recording (blue) and treatment recording (red). The bolded black traces represent the mean LFP waveform from which the duration was measured. C, Average number of LPFs detected from baseline and corresponding treatment recordings (*n* = 5). Data in red represent the mean across all slices. D and E, Average peak and valley amplitude calculated from baseline and treatment recordings. F, Average duration of LFPs recorded during baseline and corresponding treatment groups. n.s., not significant, **p<0.01; paired t-test.

To further dissern the effects of Nav1.2 inhibition on 4-AP induced hyperexcitability, we asked whether partial inhibition of Nav1.2 by the ICA cmpd would reduce activity. To address this, we additionally tested the effects of a higher concentration of the ICA cmpd (500nM) predicted to inhibit Nav1.1, Nav1.3 and also Nav1.2 by approximatly fifty percent according to the IC50 values we observed, while having little to no effect on Nav1.6. We obtained similar results to those obtained with 125nM ICA cmpd, with no significant difference in the number of ictal LFPs or peak and valley amplitues, but a signficant reduction in the duration of the LPFs (Figure 6). These data suggest that partial inhibition of Nav1.2 with the ICA cmpod does not overtly alter 4-AP induced hyper-excitability, and therefore the effects observed with the 801 cmpd are likey due to inhibition of Nav1.6.

**Figure 6.**
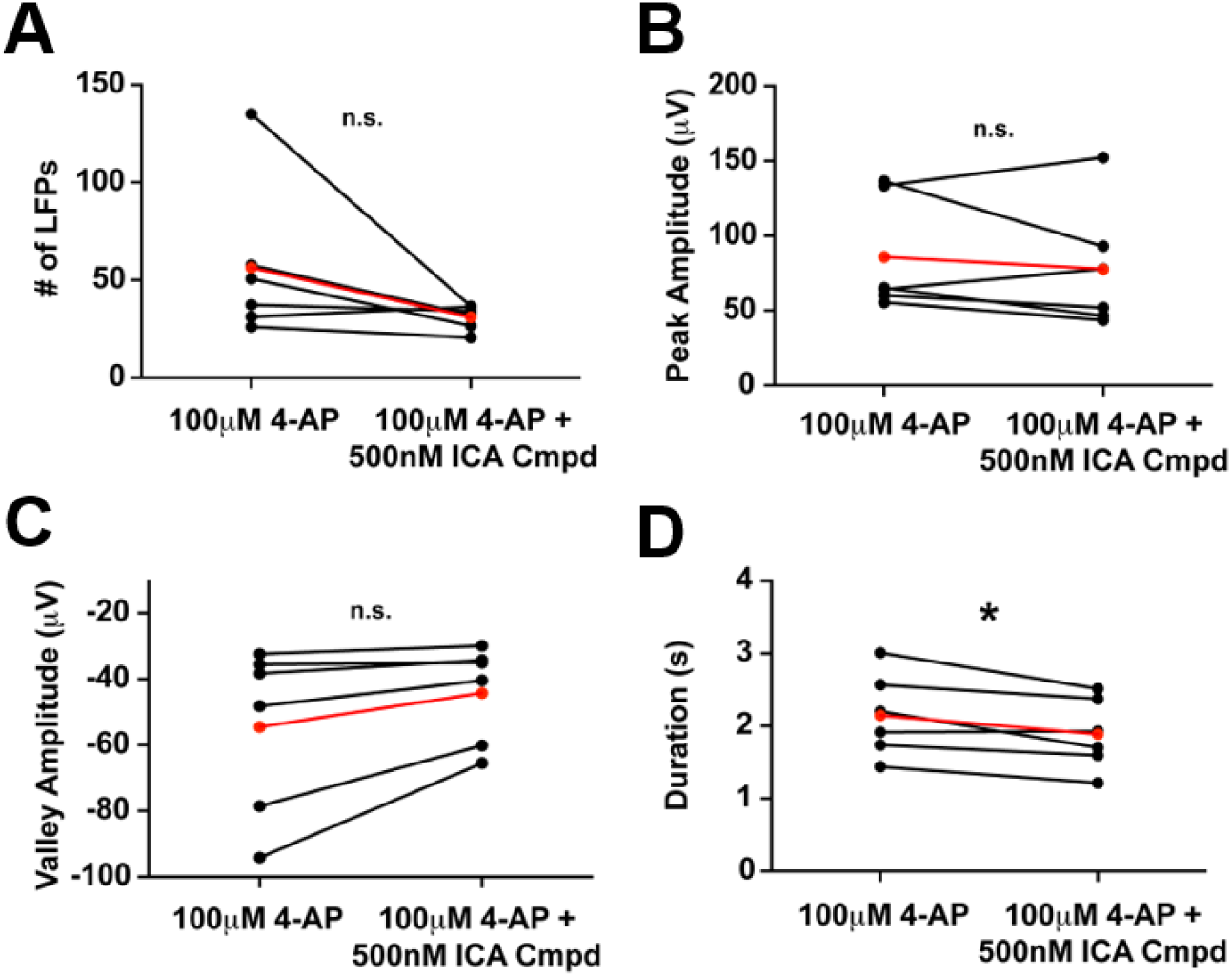
Effects of 500nM ICA compound on 4-AP induced hyperexcitability in mouse cortical brain slices. A, Number of LFPs detected during baseline (100μM 4-AP) and corresponding treatment (100μM 4-AP + 500nM ICA cmpd) recordings (*n* = 6). B and C, Average peak and valley amplitudes for LFPs recording during baseline and treatment conditions. D, Average duration of LFPs recorded during baseline and treatment recordings. C and D, n.s., not significant, *, p<0.05; paired t-test.

### Partial inhibition of Nav1.1 increases basal activity

The prominent expression of Nav1.1 in GABAergic neurons (ref) led us to ask whether partial inhibition of Nav1.1 is sufficient to induce synchronized activity. To address this question, we recorded basal activity in the presence of normal ACSF or 300nM ICA cmpd and subsequently perfused 100μM 4-AP at the end of each recording to confirm slice viability (Figure 7A). While statistical significance was not reached, we found a trend towards an increase in the number of LFPs in the presence of 300nM ICA cmpd. Moreover, the LFPs recorded in the presence of 300nM ICA cmpd were significantly larger in magnitude (green bars; *n* = 5) (p<0.05; unpaired t-test), both peak and valley amplitudes, compared to LFPs recorded in the presence of normal ACSF (black bars; *n* = 3) (Figure 7B-D). This data suggests that partial inhibition of Nav1.1 can increase basal synchronized activity.

**Figure 7.**
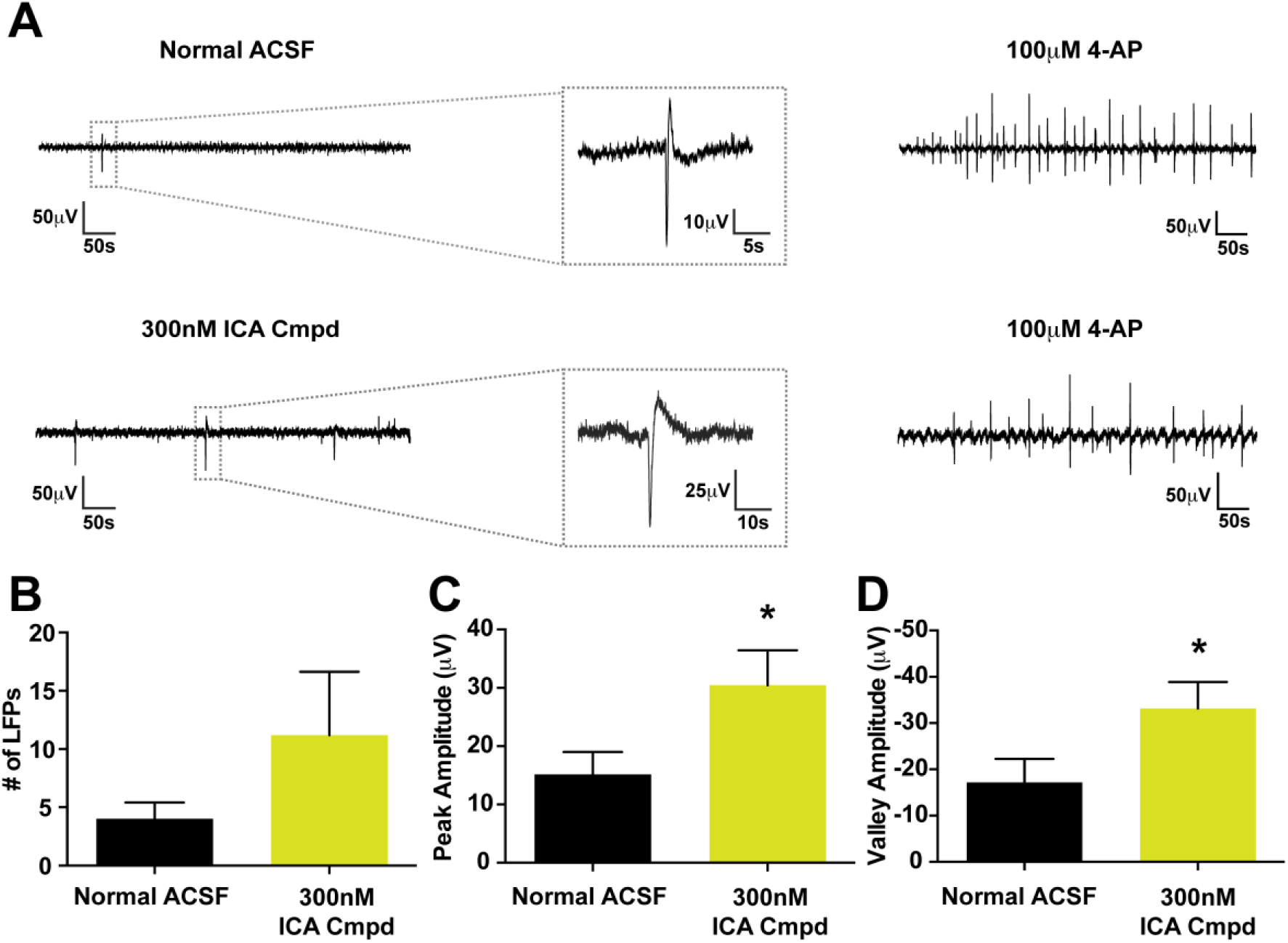
Effects of ICA compound on basal activity in mouse cortical brain slices. A, Representative twenty minute traces of basal activity from recordings in the presence of normal ACSF (top) and ACSF containing 300nM ICA cmpd (bottom). B, Average number of events detected from normal ACSF (*n* = 3; black bars) and ACSF containing 300nM ICA cmpd (*n* = 5; green bars). C and D, Average peak and valley amplitude calculated for each condition. *p<0.05; unpaired t-test.

## DISCUSSION

In this study, we examined the contribution of VGSC isoforms to epileptiform activity in order to further our understanding of the roles of specific isoforms in the brain. We tested the selectivity of recently discovered small molecule inhibitors, ICA and 801 cmpds, against Nav1.1, Nav1.2 and Nav1.6 activity in HEK293T cells. We found that the ICA cmpd is relatively selective for Nav1.1 and the 801 cmpd is a potent inhibitor of Nav1.6 and Nav1.2, which allowed us to interrogate the roles of these different isoforms. To address our main question of how specific VGSCs isoforms contribute to epileptiform activity, we observed the effects of the ICA and 801 cmpds on 4-AP induced hyper-excitability, an established *in vitro* model of epilepsy [44, 45], with MEA recordings from mouse cortical brain slices. Preferentially inhibiting Nav1.6 and Nav1.2 activity with 500nM of the 801 cmpd completely abolished ictal LFPs induced by 4-AP, but preserved slice activity. In contrast, preferentially inhibiting Nav1.1 along with Nav1.3 with 500nM of the ICA cmpd had minimal effect on 4-AP induced hyper-excitability. These findings are consistent with our current understanding of the localization and expression of these channels in the brain. Our findings clearly demonstrate that brain isoforms of VGSCs have distinct roles in epileptiform activity and suggest that selectively targeting Nav1.6 and/or Nav1.2 activity may be a more efficacious treatment strategy for epileptic syndromes than broad spectrum sodium channel inhibition.

It has been demonstrated that *Scn1a*^*+/-*^ heterozygous mice display spontaneous seizures mimicking a SMEI phenotype [7, 27]. However, it is still unclear if these seizures arise because of the direct loss of Nav1.1 or due to other compensatory mechanisms that often occur in genetic models. While some studies have found decreased somatic sodium currents of GABAergic neurons in *Scn1a*^*+/-*^ heterozygous mice [27], others report no change in somatic sodium currents but still find an impairment in excitability of GABAergic neurons [11, 50]. Therefore we also asked whether partial inhibition of Nav1.1 could alter basal slice activity due to the prominent expression of Nav1.1 in inhibitory neurons. We hypothesized that inhibition of Nav1.1 might in fact increase epileptiform activity, however we did not observe this in our preparation. We found that inhibition of Nav1.1 with the ICA cmpd, in the absence of 4-AP, increased basal synchronized activity, increasing the peak and valley amplitudes of basal LFPs along with a trend toward an increase in the overall number of basal LFPs compared to normal ACSF. It is possible that an increase in basal synchronized activity could create a circuit more susceptible to seizure generation upon a secondary insult. Indeed, a hallmark of SMEI due to *Scn1a* mutations is the initial development of febrile seizures (seizures associated with fever), which later progress into other seizure types [51].

As mentioned previously, Nav1.1 is prominently expressed in inhibitory GABAergic neurons, specifically those that express somatostatin or parvalbumin [7, 12]. Parvalbumin positive GABAergic neurons consist of two major subtypes, basket cells and chandelier cells, that provide strong synaptic inhibition to the proximal dendrites/soma and axon initial segment, respectively, of pyramidal neurons – placing them in a critical position for regulating circuit excitability [52]. Impaired excitability of parvalbumin positive neurons has been associated with epilepsy and a number of other neurological disorders [53], possibly due to their importance in the generation of gamma oscillations [54, 55]. Therefore, our data is in accordance with our current understanding of the functional role of Nav1.1 in the brain and supports the idea that perhaps Nav1.1 is not a good target for the treatment of epilepsy syndromes; indeed avoiding inhibition of Nav1.1 will likely have therapeutic benefits. Interestingly, it has recently been shown that selectively hyper-activating Nav1.1 channels in *Scn1a*^*+/-*^ heterozygous mice can reduce seizure activity and enhance survival [56].

Both Nav1.2 and Nav1.6 have been implicated in epilepsy as evidenced by genetic mutations in these channels that result in epileptic syndromes [31, 57]. Interestingly, the epilepsy-associated missense mutations in Nav1.2 have been reported to cause both gain- and loss-of function effects on channel properties when expressed in heterologous expression systems [58-60]. While this may seem paradoxical, it is not entirely surprising due to the dense expression of Nav1.2 in the proximal axon initial segment of both excitatory, pyramidal neurons as well as somatostatin positive GABAergic, inhibitory neurons. Interestingly, global inhibition of Nav1.2 results in a decreased threshold for recurrent network activity, possibly suggesting that the effects of Nav1.2 inhibition in somatostatin positive GABAergic neurons might outweigh the effects of Nav1.2 inhibition in pyramidal neurons [10].

For both the human and mouse channels, the 801 cmpd was slightly more potent at inhibiting than Nav1.2. Nav1.6 is highly expressed in the distal axon initial segment and is reportedly more important in initiation of action potentials than Nav1.2 [19]. While genetic studies support the idea that partial loss of Nav1.6 reduces seizure generation [61], they also demonstrate the potential adverse consequences that have been associated with loss of Nav1.6 activity including: absence seizures, motor defects and cognitive defects [62-65]. Genetic studies resulting in global loss of one allele of Nav1.6 may induce compensatory changes that could contribute to these adverse effects. In contrast pharmacological inhibition of Nav1.6 could possibly be titrated to achieve a balance wherein the benefits of seizure control could outweigh the consequences of unwanted side effects.

Overall, we found that partially inhibiting Nav1.6 and Nav1.2 activity pharmacologically can abolish ictal LPFs in a 4-AP model of epilepsy, while maintaining slice activity. Moreover, the compounds used in this study demonstrate voltage-dependent inhibition, similar to current AEDs that broadly target voltage-gated sodium channels [49], which allows for the selective targeting of hyper-excitable neurons, further reducing unnecessary inhibition of activity. Thus our findings suggest that selective inhibition targeting of Nav1.6 and/or Nav1.2 isoforms, rather than broadly inhibiting all VGSC isoforms as many current AEDs do, is likely to reduce unwanted side effects.

## ACKNOWLEDGEMENTS

The Compound 801 compound was generously provided by Eli Lilly and Co. (Indianapolis, IN). The research was supported by a National Institutes of Health grant (NS053422) and a grant from the Dravet Syndrome Foundation to T.R.C.. R.R.P. was partially supported by a Paul and Carole Stark Medical Neuroscience Graduate Student Fellowship.

## Notes

### Competing Interest Statement

The authors have declared no competing interest.

